# Learning of complex auditory patterns changes intrinsic and feedforward effective connectivity between Heschl’s gyrus and planum temporale

**DOI:** 10.1101/848416

**Authors:** Massimo Lumaca, Martin J. Dietz, Niels Chr. Hansen, David R. Quiroga-Martinez, Peter Vuust

**Affiliations:** Center for Music in the Brain, Department of Clinical Medicine, Aarhus University & The Royal Academy of Music Aarhus/Aalborg, Nørrebrogade 44, 8000 Aarhus, Denmark; Center of Functionally Integrative Neuroscience, Institute of Clinical Medicine, Aarhus University, Nørrebrogade 44, 8000 Aarhus, Denmark; The MARCS Institute for Brain, Behaviour, and Development, Western Sydney University, Locked Bag 1797, Penrith NSW 2751, Australia

**Keywords:** fMRI, DCM, auditory cortex, learning, predictive coding, effective connectivity, precision-weighting

## Abstract

Learning of complex auditory sequences such as language and music can be thought of as the continuous optimisation of internal predictive representations of sound-pattern regularities, driven by prediction errors. In predictive coding (PC), this occurs through changes in the intrinsic and extrinsic connectivity of the relevant cortical networks, whereby minimization of precision-weighted prediction error signals improves the accuracy of future predictions. Here, we employed Dynamic Causal Modelling (DCM) on functional magnetic resonance (fMRI) data acquired during the presentation of complex auditory patterns. In an oddball paradigm, we presented 52 volunteers (non-musicians) with isochronous 5-tone melodic patterns (standards), randomly interleaved with rare novel patterns comprising contour or pitch interval changes (deviants). Here, listeners must update their standard melodic models whenever they encounter unexpected deviant stimuli. Contour deviants induced an increased BOLD response, as compared to standards, in primary (Heschl’s gyrus, HG) and secondary auditory cortices (planum temporale, PT). Within this network, we found a left-lateralized increase in feedforward connectivity from HG to PT for deviant responses and a concomitant disinhibition within left HG. Consistent with PC, our results suggest that model updating in auditory pattern perception and learning is associated with specific changes in the excitatory feedforward connections encoding prediction errors and in the intrinsic connections that encode the precision of these errors and modulate their gain.

**Significance statement:** The learning of complex auditory stimuli such as music and speech can be thought of as the continuous optimisation of brain predictive models driven by prediction errors. Using dynamic causal modelling on fMRI data acquired during a melodic oddball paradigm, we here show that brain responses to unexpected sounds were best explained by an increase in excitation within Heschl’s gyrus and an increase in forward connections from Heschl’s gyrus to planum temporale. Our results are consistent with a predictive coding account of sensory learning, whereby prediction error responses to new sounds drive model adjustments via feedforward connections and intrinsic connections encode the confidence of these prediction errors.

## Introduction

Predictive coding (PC) is a unifying theory of brain function that formally links human learning and neuroplasticity (Friston, 2002). The key notion in PC is that the brain embodies an internal model of the environment, which constantly generates predictions on the future sensory input. When predictions fail, the brain generates ‘prediction error signals’ that are weighted by their precision and passed from lower to higher levels of the brain’s hierarchy via changes in feedforward connections, thereby revising the model’s predictions (Dietz et al., 2014). Learning can thus be thought of as a process of recurrent model updating through prediction error signaling. Specifically, this entails a release of the excitatory pyramidal cells that encode prediction error from top-down expectations, accompanied by an increase in their synaptic gain or sensitivity to ascending prediction errors (Koelsch et al., 2019). This hypothesis, however, lacks experimental evidence regarding the rich and complex stimulation that occurs in natural sound environments, such as speech and music (McDermott et al., 2013).

Melodic learning is an excellent model to address this issue. Within PC, melodic learning reflects the update of internalized probabilities acquired through statistical learning and relates to melodic continuations (Hansen and Pearce, 2014) at two pitch-processing levels: (1) ‘contour’ − the rise and fall of pitch changes− and (2) ‘interval’ − the distance between adjacent tones (Schmuckler, 2016). The maintenance and updating of melodic predictive models have been studied using auditory oddball paradigms, whereby magnetoencephalographic and/or electrophysiological (M/EEG) brain activity is recorded while participants listen to streams of repeated short melodies typically based on tones from the 12-tone-equal-tempered scale, omnipresent in Western tonal music (e.g., Tervaniemi et al., 1994; Hsu et al., 2015; Lappe et al., 2016). The mismatch negativity (MMN) elicited by occasional contour or interval deviations in these patterns is evidence that a melodic predictive model has been formed during the auditory stimulation and thought to represent the message passing of precision-weighted prediction errors in the cortical auditory hierarchy (Vuust et al., 2009, 2014, 2018; Dietz et al., 2014; Auksztulewicz and Friston, 2015). Previous functional magnetic resonance imaging (fMRI) work has found core regions of the superior temporal gyrus (STG), including primary auditory cortex (A1) and planum temporale (PT), as the main cortical generators of melodic MMN (Habermeyer et al., 2009). However, how auditory cortical regions interact to form and revise melodic representations remains an unsettled issue.

Here, we used dynamic causal modelling (DCM) of effective connectivity in auditory fMRI data acquired while 52 human volunteers (non-musicians) were presented with an unfamiliar musical tuning system, the Bohlen Pierce (BP) scale (Mathews et al., 1988) (Fig. 1A). This artificial scale has the advantage of preserving intervallic properties from the 12-tone equal-tempered scale without being contaminated by prior knowledge about pitch categories and intervals. This ensures a more controlled setting to test hypotheses about the generation and revision of an internal predictive model. First, we identified an MMN auditory network using a melodic oddball paradigm (Lumaca and Baggio, 2016). In a single run, participants were scanned while listening to streams of five-tone BP melodic patterns (Fig. 1B). Occasionally, the fourth tone was pitch-shifted to violate the contour (‘contour’ deviant; 10%) or the interval structure (‘interval’ deviant; 10%) of the ‘standard’ sequence. Second, we used a two-state DCM (Marreiros et al., 2008) to model the effective connectivity within and between bilateral primary and secondary auditory cortices during deviant stimuli, relative to standard stimuli (Fig. 2). Using Bayesian model reduction (BMR) and parametric empirical Bayes (Friston et al., 2016), we then compared, at the group level, a series of alternative hypotheses (Fig. 3) to examine the specific changes in extrinsic and intrinsic effective connectivity that occur in the localized temporal network during auditory pattern learning.

**Fig. 1.**
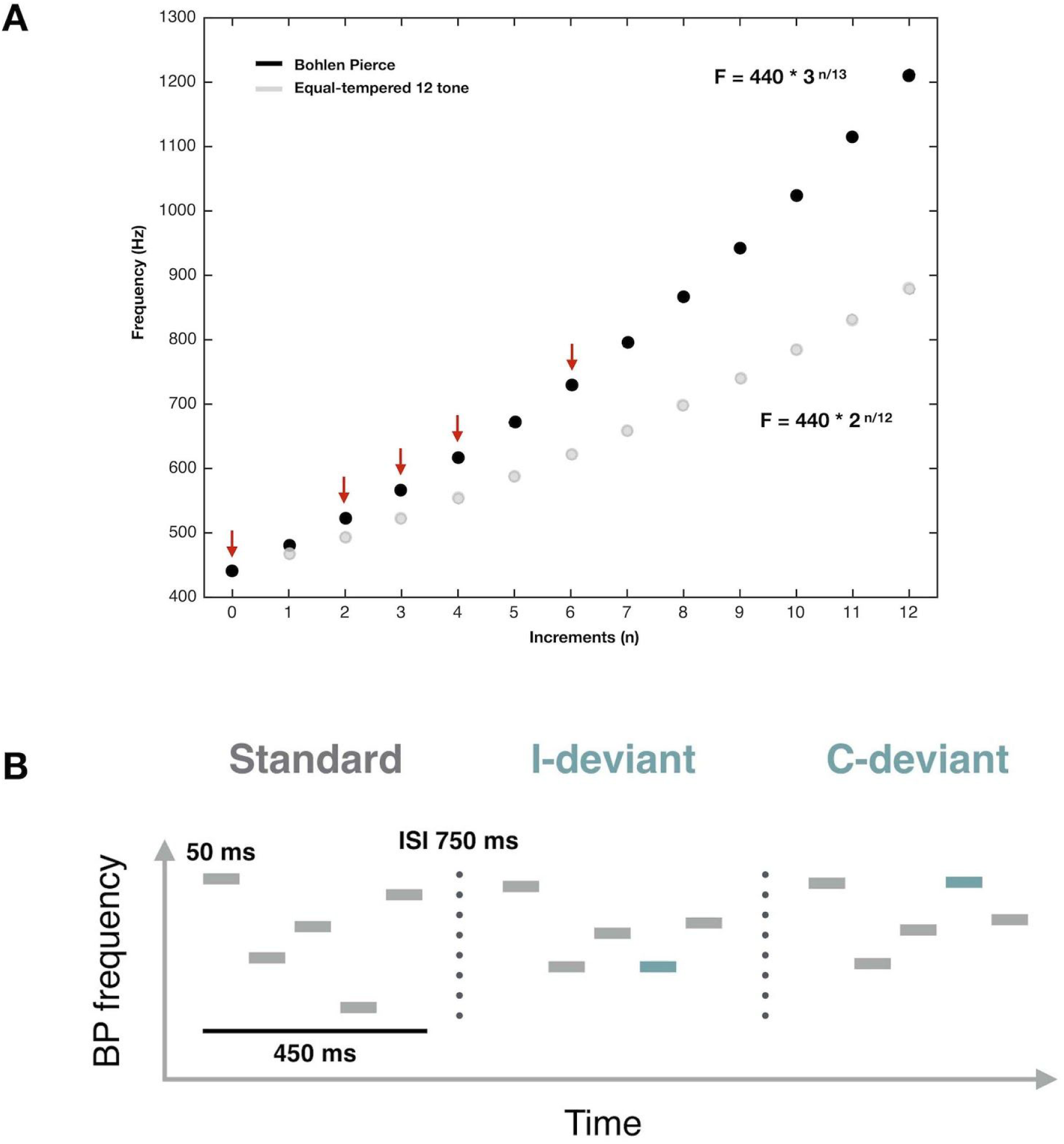
Musical scale and auditory patterns used in the current study. (A) Illustration of pitch frequencies along the Western equal-tempered 12-tone scale (in grey) and the Bohlen Pierce scale (in black). Red arrows point to the BP frequencies used to build melodic material (*k* = 440 Hz; n=0, 2, 3, 4, or 6). (B) Schematic illustration of the melodic patterns presented to participants during the auditory oddball paradigm (adapted from (Lumaca et al., 2019)). Participants were scanned while listening to these melodic patterns. Each pattern was 450-ms long, and consisted of five 50-ms sinusoidal tones separated by 50-ms silent intervals. Melodic patterns were presented with 750-ms of interstimulus interval (ISI) randomly at three frequency levels (lowest frequency: 440, 478, 567 Hz) belonging to the Bohlen Pierce musical scale. Standard patterns (80%) followed the abstract rule EBCAD. In deviant patterns, the fourth tone was changed in frequency compared to its standard position, either producing a change in the melodic interval (I-deviants; 10%) or in the melodic contour (C-deviant; 10%).

**Fig. 2.**
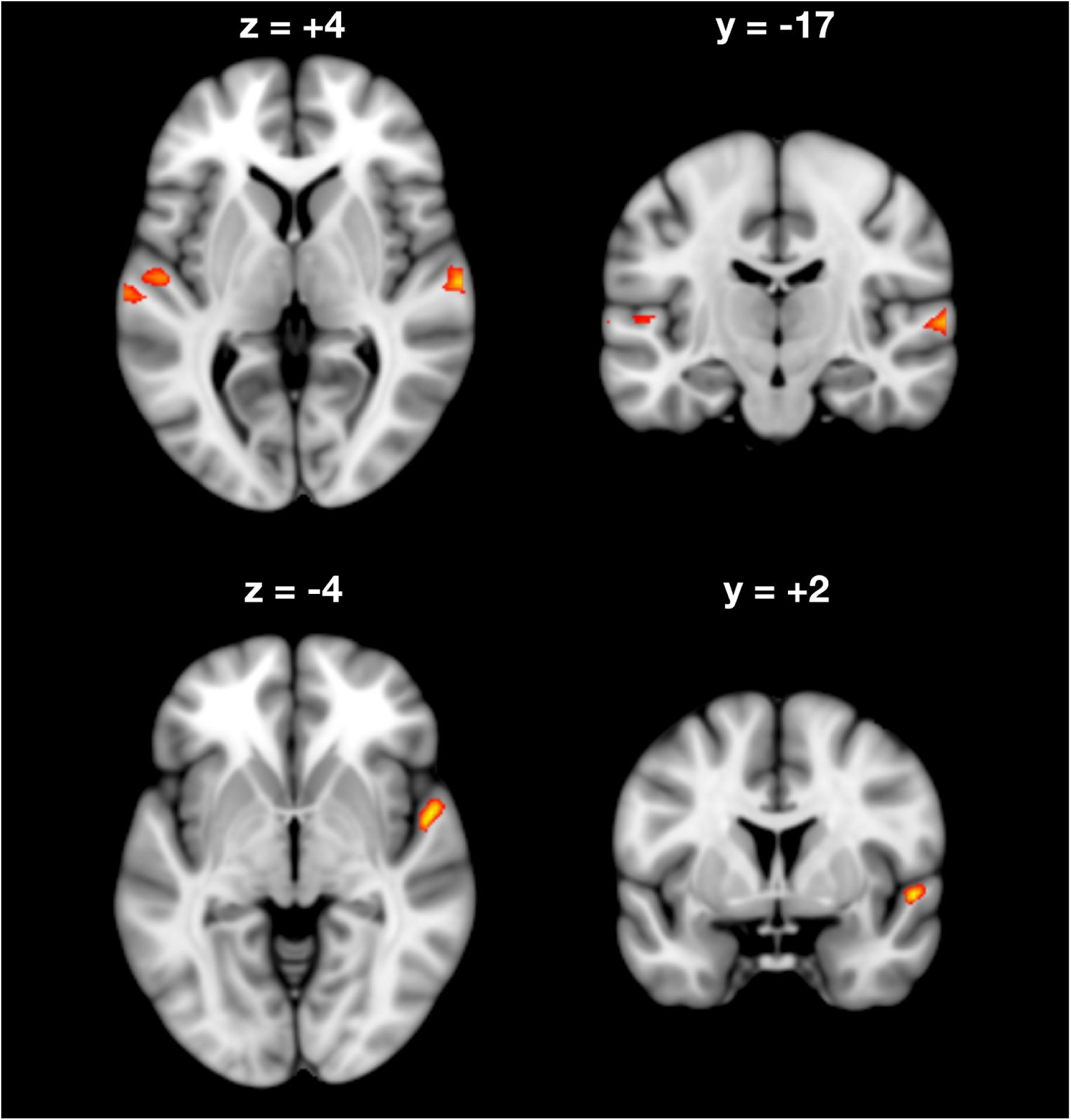
Cortical areas responding to contour deviants in melodic patterns. Shown in the figure are the cortical loci where event-related activity was greater for contour deviant events (C-deviants) compared to the standard events occurring in the same position of the pattern (i.e., the 4th tone) (*P* < 0.001, FWE corrected). Activated areas are shown projected onto an MNI standard template, and include bilateral Heschl’s Gyrus (HG) and planum temporale (PT).

**Fig 3.**
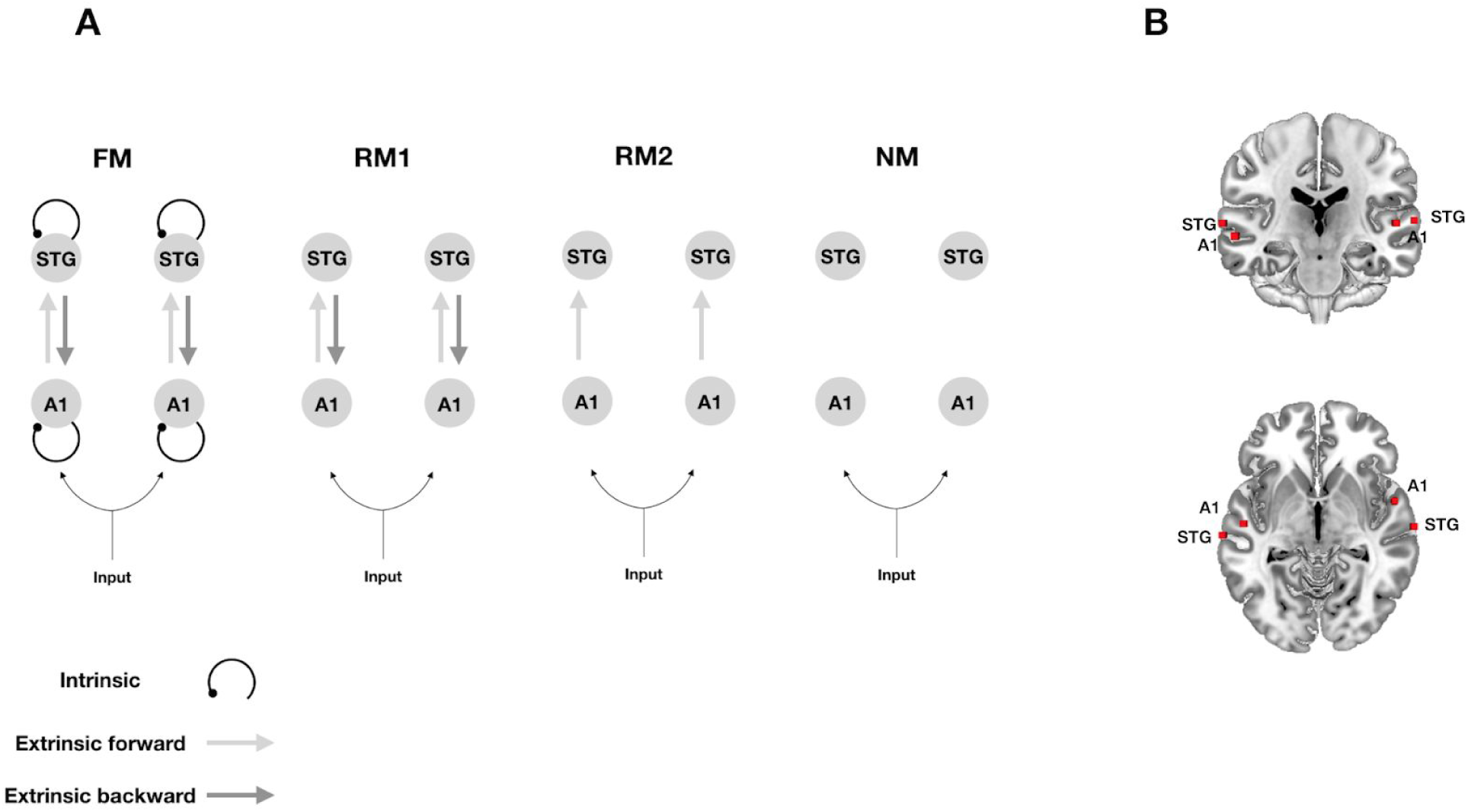
Alternative hypotheses about effective connectivity (A) The dynamic causal models comprise a bilateral ‘input’, four ‘sources’ and ipsilateral ‘connections’ among (extrinsic connections) and within (intrinsic connections) these sources. A1, primary auditory cortex; STG, superior temporal gyrus. The four models tested had the same anatomical architecture but differed in terms of the embedding connections: intrinsic and extrinsic (forward and backward) in the full model (FM), only forward and backward in the reduced model 1 (RM1), only forward in the reduced model 2 (RM2), and no connections in the null model (NM). (B) Sources (red squares) were defined by using peak activations of the auditory oddball localiser, and are here projected onto an anatomical MNI standard template.

## Materials and Methods

### Participants

A total of 52 participants (33 females, mean age 24.5 years, range 20-34) with normal hearing took part in the fMRI experiment. Participants were all non-musicians (i.e., none of them had three or more years of formal musical training) and all gave informed consent before the experiment. The neuroimaging data used in this work were acquired a part of another project approved by the local ethics committee of the Central Denmark Region (nr. 1083).

### MRI Procedure

#### Bohlen-Pierce scale

The auditory stimulation used in our fMRI oddball paradigm was adopted from an EEG paradigm by Lumaca and Baggio (2016). Tone sequences were constructed using tones from the equal-tempered version of the Bohlen-Pierce (BP) scale (Mathews et al., 1988; Loui et al., 2009), a microtonal tuning system with 13 logarithmic even divisions of a tritave (corresponding to a 3:1 frequency ratio). In the equal-tempered version of the scale, frequencies (F) are defined by the following:

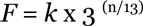

where *n* is the number of steps along the scale, and *k* is a constant that correspond to the fundamental frequency. Based on this equation, we defined n = 0, 2, 3, 4 or 6 and *k =* 440 Hz (Fig. 1A).

#### Stimuli

The pattern deviance paradigm and stimuli were adapted from an EEG paradigm by Lumaca and Baggio (2016), and has been shown to produce a MMN evoked potential (Näätänen et al., 1978). Stimuli were 5-tone melodic patterns presented in a single block, consisting of collections of five 50 ms sinusoidal tones (5 ms rise and fall; 50 ms inter-tone intervals) with the frequencies 440, 521, 567, 617, 730.6 Hz (henceforth referred to as ABCDE in the context of the low register) (Fig. 1B). During stimulation, sequences were randomly transposed to three different baseline frequencies (henceforth referred as ‘A’), corresponding to three different registers of the BP scale (baseline tones: 440, 478, 567 Hz). The patterns were presented with an ‘inter-stimulus interval’ of 750 ms. Standard patterns (80%) were randomly interspersed with ‘contour’ (10%) and ‘interval’ (10%) deviant sequences for a total of 1260 stimuli. In abstract terms, the standard pattern followed the sequence EBCAD. In contour deviant stimuli, the fourth tone changed the surface structure (‘ups’ and ‘downs’) of the standard stimuli without changing the interval (i.e. EBCED); viceversa for the interval deviants (i.e. EBCBD). Deviant stimuli were pseudo-randomized in order to present deviant tones of the same type in close temporal succession (jittering stimulus-onset asynchrony, SOA, range 2400-4800 ms) and to induce an increase of the BOLD response by superposition. The use of multiple deviants in close succession can be found in other fMRI oddball studies (Cacciaglia et al., 2015).

#### Image acquisition

The fMRI data were acquired on a 3T MRI scanner (Siemens Prisma). The subjects’ head was fixated with cushions to minimize movement during the experiment. During MRI acquisition, participants were instructed to be still and to watch a subtitled silent movie projected on a MRI-compatible screen located at the rear of the scanner. In the meanwhile, auditory stimulation was delivered by MR-compatible headphones using Presentation software (www.neurobs.com). A total of 1535 volumes were acquired over 25 minutes using a fast T2*-weighted echo-planar imaging (EPI) multiband sequence (TR, 1000 ms; TE, 29.6 ms; voxel size, 2.5 mm^3^). A T1-high resolution image was also acquired using an MP2RAGE sequence (TR, 5000 ms; TE=2.87 ms; voxel size, 0.9 mm^3^).

#### Preprocessing and analyses

Image time-series were preprocessed and analyzed using SPM12 (r7487), implemented in Matlab R2016b (Mathworks). EPI images were first spatially realigned to the first EPI volume. Then, individual high-resolution T1-images were coregistered to the mean EPI image and segmented using the standard tissue probability maps of SPM12. All functional EPI images were then normalized to a standard Montreal Neurological Institute (MNI) reference brain using the resulting deformation fields, resliced to 2 × 2 × 2 mm^3^ and smoothed with an isotropic 6 mm Gaussian kernel. Low frequency noise was removed through the use of a high-pass filter (cutoff 1/128 Hz), and time-series were corrected for serial autocorrelations using a first-order autoregressive (AR(1)) model.

First-level analysis consisted of a general linear model (GLM) with standard (STD; implicitly modelled), deviant contour (C-deviant), and deviant interval (I-deviant) regressors convolved with a canonical HRF, plus realignment parameters to account for head motion. At the second level, we used a whole-brain random-effects analysis using t-test contrasts C-deviant>STD and I-deviant>STD at *p*<0.001 familywise error (FWE) corrected at voxel-level.

#### Volume-of-interest (VOI) extraction

We summarised the BOLD signal in each participant using the first eigenvariate (principal component) of voxels within a sphere of 8 mm radius centred on each participant’s local maximum. This subject-specific local maximum was identified within a sphere of 20 mm radius centred on the peak of the group effect.

#### Dynamic causal modelling of effective connectivity

We used a two-state dynamic causal model (DCM) for fMRI (DCM12, revision 7487) to estimate the effective connectivity between Heschl’s gyrus (HG) and Planum Temporale (PT) within each hemisphere as well as the intrinsic connectivity within each of these regions, given observed haemodynamic measurements (Friston et al., 2003). In two-state DCM, each region comprises one excitatory and one inhibitory population of neurons. This allows us to model the intrinsic connectivity within each cortical area as an increase or decrease in cortical inhibition (Marreiros et al., 2008).

Although fMRI is an indirect measure of neuronal activity based on the observed blood oxygenation level dependent (BOLD) signal, the biophysical model employed in DCM is equipped with a detailed haemodynamic forward model that describes how neuronal activity translates into changes in regional blood flow (Friston et al., 2000), blood volume and deoxyhemoglobin concentration (Buxton et al., 1998) that combine non-linearly to produce the BOLD signal (Stephan et al., 2007). This means that the neuronal model comprised of excitatory and inhibitory populations can be used to make inferences about both long-range excitatory connectivity between brain regions as well as local connectivity within a region that reflects changes in the ratio of excitatory and inhibitory activity (Logothetis, 2008).

Hemodynamic responses to all auditory stimuli (standard stimuli and contour deviants) were modelled as a driving input to HG in both hemispheres. Using parametric modulation of the regressor encoding all stimuli, responses to contour deviants compared to standard stimuli were then modelled as a modulatory input to the network under four alternative hypotheses about how connection strengths change during auditory contour deviancy: The first DCM comprised a full network, where changes in both feedforward and feedback connections between HG and STG, as well as their intrinsic (inhibitory) connections, encode the differences between deviant and standard conditions. Within predictive coding, this hypothesis represents the belief that both forward prediction errors and backward predictions mediate the statistical learning of melodic regularities and that the intrinsic connectivity may encode the precision with which prediction error are broadcasted during belief updating. The second DCM was formulated as a reduced model where only feedforward and feedback connections between HG and STG encode the differences between deviant and standard conditions. This hypothesis represents the belief that both prediction errors and predictions mediate the statistical learning of melodic regularities, in the absence of notable changes in the intrinsic connectivity encoding the precision. The third hypothesis was formulated as another reduced model where only the forward connections from HG to STG encodes the differences between deviant and standard conditions. This hypothesis represents the belief that only forward prediction errors mediate the statistical learning of melodic regularities, in the absence of notable changes in feedback and intrinsic connections. The fourth hypothesis was a null model, encoding the belief that no cortical connections change during statistical learning of melodic regularities (see Figure 3A for a schematic of alternative hypotheses). We then inverted the full model using Variational Laplace (Friston et al., 2007). This provides the posterior probability of connection strengths and the free-energy approximation to the Bayesian model evidence. The reduced models and the null model were then estimated using Bayesian model reduction (Friston et al., 2016).

#### Parametric Empirical Bayesian (PEB) analysis of group effects

We then used parametric empirical Bayes (PEB) (Friston et al., 2016; Zeidman et al., 2019) to identify increases or decreases in extrinsic (excitatory) connections between HG and STG and intrinsic (inhibitory) connections within each region at the group level. PEB is a hierarchical Bayesian model in which empirical priors on the connection strengths at the first (single-subject) level are estimated empirically from the data themselves using a Bayesian general linear model (GLM) at the group level. In this way, PEB provides both the posterior probability of connection strengths and the Bayesian model evidence for Bayesian inference and model comparison at the group level. The model evidence of the PEB model is given by the sum of DCM accuracies for all participants, minus the complexity of both the first-level DCMs and the second-level Bayesian GLM. During PEB estimation, we used the updated DCM parameters where the connection strengths had been re-evaluated using the group means as priors to obtain the most robust estimates (Zeidman et al., 2019). An advantage of PEB, as opposed to classical random-effects (RFX) analysis, is that PEB takes not only the mean, but also the uncertainty of individual connection strengths into account. This means that participants with more uncertain parameter estimates will be down-weighted, while participants with more precise estimates receive greater influence (Zeidman et al., 2019).

## Results

### Auditory-cortex functional localizer

Contour deviant stimuli produced significantly greater activation than standard stimuli in bilateral STG (Table 1). Specifically, the contrast C-deviant > STD revealed four clusters of activation (*p*_FWE_ < 0.001): two localized in the left and right Heschl’s Gyrus (HG; cytoarchitectonic areas Te 1 and Te 1.2) and two more posterior localized in the planum temporale (PT; cytoarchitectonic area Te 3) (Morosan et al., 2005) (Fig. 2). Conversely, the contrast between interval deviant stimuli and standard stimuli (I-deviant > STD) did not produce any significant activation (*p*_FWE_ > 0.05).

**Table 1.** MNI coordinates of brain regions activated in the C-deviant > STD contrast (Height threshold: T = 6.76, p_FWE_ <0.001; Extent threshold: k = 0 voxels).

| *T statistic* | MNI coordinate | Anatomical region | Probabilistic atlas<sup>1</sup> |
| --- | --- | --- | --- |
| 8.83 | [54 2 -4] | Right Superior Temporal Gyrus | Area TE (1.2) 28%<br>OP4 (PV) 17% |
| 8.54 | [66 -16 4] | Right Superior Temporal Gyrus | Area TE (3) 57% |
| 7.93 | [-52 -14 4] | Left Superior Temporal Gyrus | Area TE (1) 46%<br>Area TE (1.2) 13% |
| 7.88 | [-66 -22 6] | Left Superior Temporal Gyrus | Area TE (3) 73% |

**Notes:** Negative coordinates indicate brain regions in the left hemisphere. C-deviant = contour deviant; r = right hemisphere; l = left hemisphere. <sup>1</sup>Anatomical classification using the SPM anatomy toolbox (Eickhoff et al., 2005).

### Dynamic causal modelling of effective connectivity in the auditory system

The coordinates of the four peak activations observed for the contrast C-deviant > STD were used to define volumes of interest (VOIs) in the DCM analysis (Fig. 3B). Bayesian model comparison (Fig. 3A) showed high evidence for a full DCM, where the intrinsic (inhibitory) coupling within regions and the extrinsic (excitatory) coupling between regions were modulated during deviant stimuli, relative to frequent standard stimuli (Model posterior probability > 0.99). Within this bilateral network, we observed a left-hemispheric increase in (excitatory) feedforward connectivity from HG to STG (Posterior probability > 0.99) and a concomitant decrease in the intrinsic (inhibitory) connectivity within left HG (Posterior probability > 0.99) during deviant melodic stimuli (Fig. 4). See Figure 5 for a schematic of the two-state dynamic causal model used in this study.

**Fig 4.**
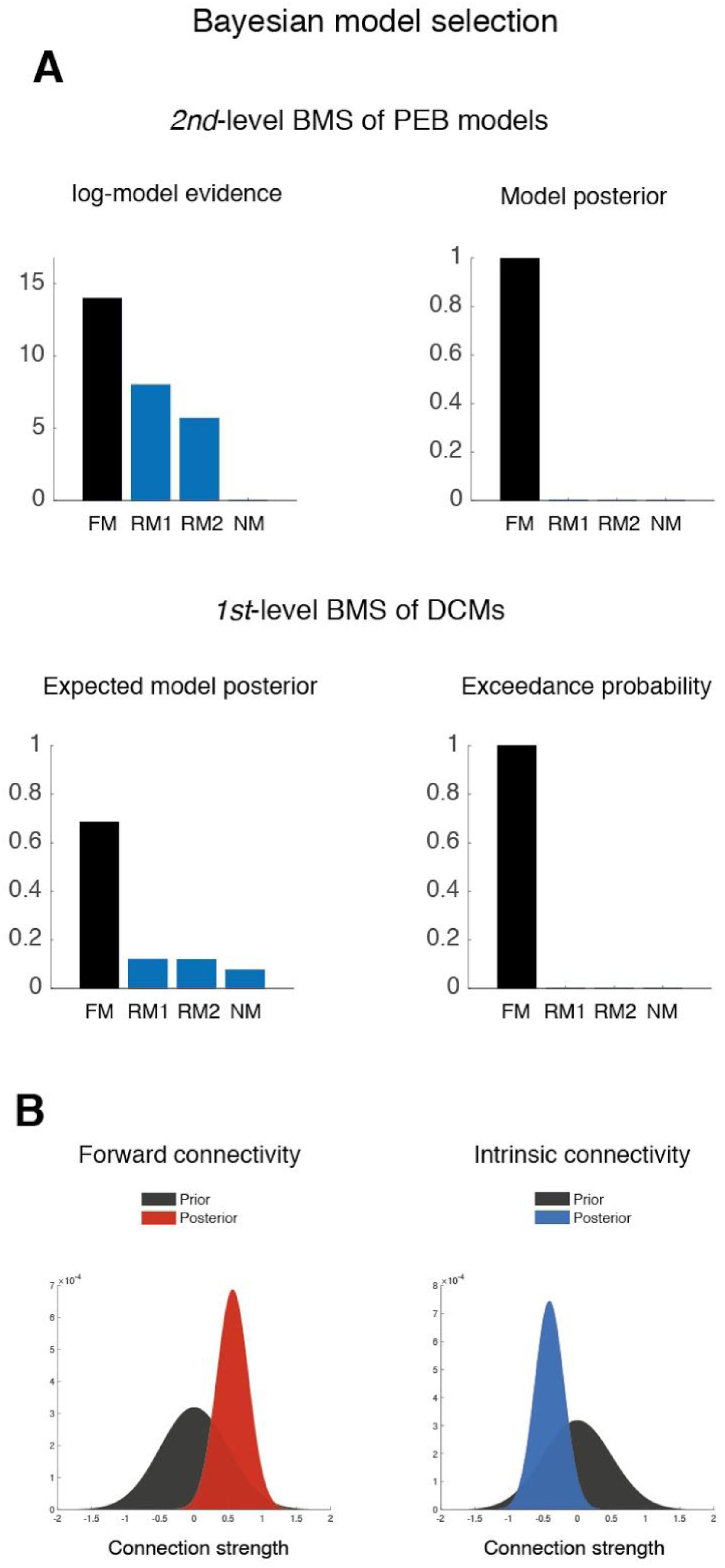
(A) Bayesian model selection of alternative hypotheses revealed that the full model with extrinsic and intrinsic connections outperforms the other models, both at the first- and the second levels of model inference. (B) Posterior probabilities of the excitatory feedforward connection strength from left HG to PT and the intrinsic inhibitory connection strength within left HG. This shows the posterior distribution of the increase in feedforward connection strength (red probability density) and the posterior distribution of the decrease in self-inhibition (blue density) as they moved away from their prior distribution (black density) after model inversion.

**Figure 5.**
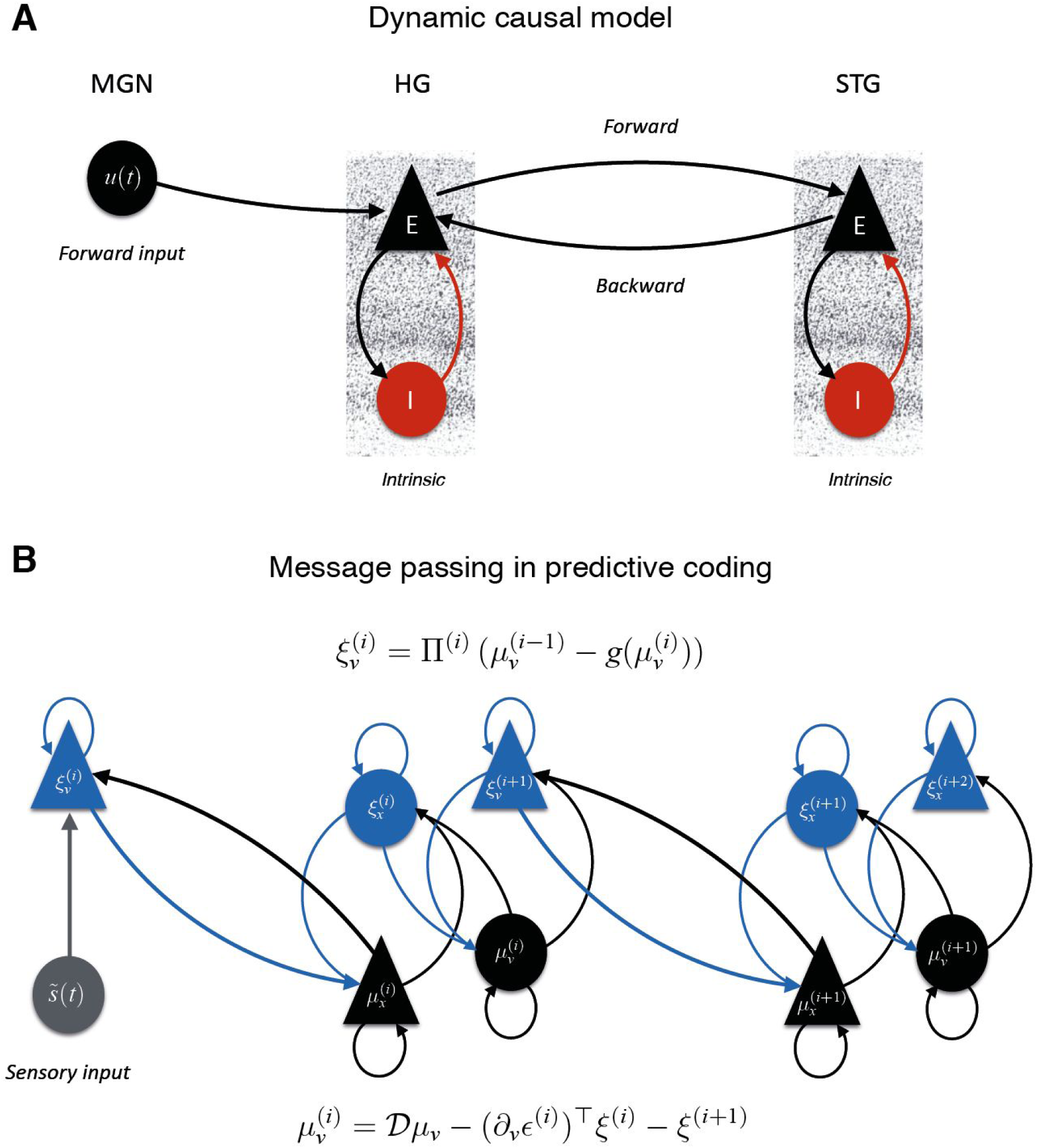
(A) Representation of the two-state DCM with one excitatory (E) and one inhibitory (I) population of neurons. In this cortical network, Heschl’s Gyrus (HG) receives input from the medial geniculate nucleus (MGN) of the thalamus and is connected to the planum temporale (PT) through (excitatory) forward and backward connections. (B) Message passing scheme proposed by predictive coding. Prediction units (in black) encode expectations (μ) about hidden causes (*v)* and hidden states (*x)*. The gain of prediction error (ξ) units (in blue) is modulated by their expected precision (Π). Superindices indicate the level of processing in the hierarchy. Time-dependent sensory input (*s(t)*) is indicated in grey.

## Discussion

Comparing different DCM models in fMRI oddball data, we found BOLD responses for deviant sounds in melodies to be best explained by a fully connected bilateral auditory network, with a decrease in inhibitory connectivity within left HG and an increase in feedforward connectivity from left HG to PT. This is consistent with a hypothesis of PC regarding sensory learning, whereby prediction errors to regularity violations are produced by an increase in synaptic gain in neuronal error units and passed forward to higher hierarchical levels. Our findings provide evidence for a generalized neuronal mechanism underlying unexpected or surprising stimuli during melodic pattern encoding.

These findings can be interpreted in the light of predictive coding (PC), which is a framework for understanding the computational mechanisms of perception and learning in the brain. The key notion in PC is that the brain creates a hierarchical generative model of its environment. Here, higher levels provide predictions or expectations about the hidden causes of sensory inputs. These causes are hidden in the sense that they are not directly observed, but can only be inferred from sensory data, given a generative model of how they were caused. When a sensory input does not conform to prior expectations, the ensuing prediction errors generated at lower levels serve to update beliefs at higher levels to optimize predictions (Friston and Kiebel, 2009).

Crucially, the relative influence of prior expectations and prediction errors on perceptual inference is controlled by their relative precision or confidence (i.e., inverse variance). This means the brain has a first-order system for expectations and prediction errors that encodes hidden causes in terms of their first-order statistics, and a second-order system that encodes the precision of first-order expectations and prediction errors in terms of their expected precision and ensuing prediction errors on the precision. Biologically, the hierarchical architecture of predictive processing is likely implemented in the brain via feedback and feedforward connections that mediate prediction and prediction errors (Bastos et al., 2012) (Fig. 5).

Our DCM results can be mapped onto the above-mentioned predictive coding scheme. Bayesian model selection shows that melodic deviance processing occurs throughout a hierarchy of bilateral superior temporal regions, with a left-hemispheric lateralization in the modulation of connection strengths during melodic deviance processing. Specifically, we provide strong evidence that mismatch responses were generated by a decrease in intrinsic (inhibitory) connectivity within left HG and an increase in feedforward (excitatory) connectivity from left HG to PT. From a predictive coding perspective, an increase in feedforward connectivity corresponds to the passing of prediction error from lower to higher areas of the hierarchical processing network, so that it can effectively update the internal generative model (Wacongne et al., 2012; Lieder et al., 2013). Similarly, the decrease in intrinsic (inhibitory) connectivity in HG can be interpreted as a precision-related increase in the gain of the superficial pyramidal cells encoding prediction error (Kiebel et al., 2007; Feldman and Friston, 2010) (Fig. 5).

In two-state DCM, this gain modulation arises from the excitation/inhibition balance of excitatory pyramidal cells and inhibitory interneurons. This interpretation entails that, besides generating prediction error signals, melodic deviants are afforded a higher precision than standards, which is consistent with the role of salient stimuli in the orientation of attention (Polich and Criado, 2006; Hannon and Trainor, 2007; Parr and Friston, 2017). Thus, unexpected sounds would point to the sources in the auditory scene that are most informative and relevant and need to be prioritized for processing through gain mechanisms. This interpretation is also in agreement with studies showing an attentional enhancement of auditory prediction error responses (Garrido et al., 2018), which has been associated with an intrinsic gain modulation in superficial pyramidal cells, mediated by a decrease in the input of inhibitory interneurons (Auksztulewicz and Friston, 2015).

Earlier DCM work on the auditory MMN using M/EEG data (e.g., Garrido et al., 2007, 2009; Kiebel et al., 2007; Dietz et al., 2014) found that temporal (HG and PT) and frontal sources (inferior frontal gyrus, IFG) in the DCM models, with forward, backward and intrinsic modulations, were the best to explain MMN evoked responses to frequency deviants. Our work is generally consistent with these findings. However, unlike previous studies, we did not find a modulation of backward connections, which may be due to a number of reasons. First, we have taken advantage of the recently developed parametric empirical Bayes (PEB) approach (Friston et al., 2016; Zeidman et al., 2019), which allows more precise inferences at the single parameter level, informed by empirical priors taken from group-level estimates, as compared to classical random-fixed effect modelling. Second, the stimulation used in past studies is inherently different from ours. In past DCM studies, participants were presented with ‘classical’ oddball stimulation, whereby trains of standard tones were randomly interleaved with frequency deviants. The more complex stimulation of the present study might have affected the connectivity of the auditory temporal network differently. Relatedly, while typical oddball stimulation has been useful to assess expectations for individual pitches, it is not adequate to investigate statistical learning of complex stimuli such as music. To our knowledge, our study is the first to look at the neural dynamics underlying the violation of complex regularities during listening to melodic patterns.

Another reason for the discrepancies might be that the neural ‘architecture’ employed in past DCM models included approximate locations of the MMN generators, which may have reduced their *construct validity* (Friston, 2003). At present, there has been a lack of anatomical accuracy in the characterization of cortical networks recruited during deviance detection. In most DCM studies, cortical sources are defined *a priori*, based on previous fMRI studies using different experimental setups and stimuli (e.g., see Garrido et al., 2007, 2009; Kiebel et al., 2007). This may have led to the inclusion of sources, such as the right inferior frontal gyrus, that fit the measured MMN waveform but that were not actual generators for the sample at hand. In some M/EEG studies, this issue was addressed with source reconstruction (Auksztulewicz and Friston, 2015; Fardo et al., 2017). However, the lower spatial resolution and inherent spatial inverse problem of M/EEG techniques, relative to fMRI, make it harder to resolve MMN cortical generators from neighboring neural populations, thus producing coarser maps at which these computational units operate. To our knowledge, this study is the first to use DCM on auditory networks functionally defined within the same fMRI sample in an event-related design.

The fMRI oddball localizer in the present study indicates that melodic contour mismatch responses are encoded in two core regions of the bilateral superior temporal plane: Heschl’s gyrus (HG) and planum temporale (PT). No significant modulation of brain activity was observed for interval changes. Contour processing is more fundamental and basic than interval processing. It is critical in the perception of music as well as speech (Patel et al., 1998), develops earlier in the ontogeny of individuals (Lamont, 2016), and changes in its content are more easily detected than interval changes (Edworthy, 1985). Conversely, encoding of interval information requires more intensive training and long retention intervals (Dowling and Bartlett, 1981). In the current experiment, the use of the Bohlen-Pierce scale ensures that there is no contamination of prior knowledge from earlier music life exposure (Ross and Hansen, 2016). The fact that a tonal hierarchy is not readily available to the participants may have hindered the detection of interval changes. It is thus not surprising that the modulation of responses to interval changes was not strong enough to produce detectable effects.

The auditory network identified in the current experiment is consistent with previous fMRI studies addressing the location of MMN generators for pitch deviants, whether using continuous oddball stimulation (e.g., Opitz et al., 2002; Liebenthal et al., 2003; Molholm et al., 2005; Schönwiesner et al., 2007) or more complex auditory patterns (Habermeyer et al., 2009). All of them reported a major activity at bilateral locations of the STG to pitch deviants, including primary and secondary auditory cortices, while only a minority reported activation of frontal areas (Deouell, 2007; Alho et al., 2014). In line with these results, we did not observe modulation of frontal lobe activity. This may be related to a putative lower signal-to-noise ratio in frontal sources compared to temporal sources, coupled with the low temporal resolution of fMRI techniques (Deouell, 2007). An alternative account is the virtual absence in our paradigm of top-down ‘schematic’ expectations, which are thought to be related to frontal areas (Koelsch, 2002; Garza Villarreal et al., 2011).

In conclusion, consistent with PC, our results show that learning of complex auditory patterns is associated with changes in excitatory feedforward connections for encoding prediction errors and changes in intrinsic connectivity for encoding the precision of prediction errors within the auditory system. Contrary to previous literature, our findings support an interpretation whereby intrinsic and extrinsic neuronal circuitry in the superior temporal gyrus ‘alone’ may establish models of short-term statistical regularities, generate predictions and update the internal model when predictions are not fulfilled.

## Author contributions

ML conceived and designed the experiment. ML acquired the fMRI data. ML preprocessed and analysed the fMRI data. MJD performed DCM analysis and Bayesian analysis. ML, MJD, and DQ prepared the figures. ML, MJD, NCH, and DQ wrote the paper. ML, MJD, NCH, DQ and PV revised and approved the final version of the manuscript.

## Acknowledgments

The authors thank Hella Kastbjerg for proofreading the manuscript, and Claudia Iorio and Ulrika Varankaite for assistance in data acquisition. Center for Music in the Brain is funded by the Danish National Research Foundation (DNRF117). MJD is funded by VELUX FONDEN (00013930). NCH is funded by Carlsberg Foundation (CF18-0668) and Lundbeck Foundation (R266-2017-3339).

